# Genetic and cellular sensitivity of *Caenorhabditis elegans* to the chemotherapeutic agent cisplatin

**DOI:** 10.1101/233023

**Authors:** Francisco Javier García-Rodríguez, Carmen Martínez-Fernández, David Brena, Ernest Nadal, Mike Boxem, Sebastian Honnen, Antonio Miranda-Vizuete, Alberto Villanueva, Julián Cerón

## Abstract

Cisplatin and derivatives are commonly used as chemotherapeutic agents. Although the cytotoxic action of cisplatin on cancer cells is very efficient, clinical oncologists need to deal with two major difficulties: (i) the onset of resistance to the drug, and (ii) the cytotoxic effect in patients. Here we use *Caenorhabditis elegans* to investigate factors influencing the response to cisplatin in multicellular organisms. In this hermaphroditic model organism, we observed that sperm failure is a major cause in cisplatin-induced infertility. RNA-seq data indicate that cisplatin triggers a systemic stress response in which DAF-16/FOXO and SKN-1/Nrf2, two conserved transcription factors, are key regulators. We determined that inhibition of the DNA-damage induced apoptotic pathway does not confer cisplatin protection to the animal. However, mutants for the pro-apoptotic BH3-only gene *ced-13* are sensitive to cisplatin, suggesting a protective role of the intrinsic apoptotic pathway. Finally, we demonstrate that our system can also be used to identify mutations providing resistance to cisplatin and therefore potential biomarkers of innate cisplatin-refractory patients. We show that mutants for the redox regulator *trxr-1*, ortholog of the mammalian Thioredoxin-Reductase-1 TrxR1, display cisplatin resistance and that such resistance relies on a single selenocysteine residue.

## INTRODUCTION

The FDA approved the use of cisplatin (CDDP, *Cis-DiammineDichloride Platinum* (ii)) as chemotherapeutic agent in 1978. Since then, cisplatin and other platinum-based derivatives have successfully been used in cancer treatment. To illustrate their impact in the clinic, it has been estimated that approximately half of all patients undergoing chemotherapeutic treatment receive a platinum drug ^1^. Cisplatin exerts activity against a wide spectrum of solid neoplasms, including testicular, bladder, ovarian, head and neck, gastric or lung cancers ^2^. Strikingly, testicular cancer was previously fatal but treatment with cisplatin provided cure for 80% of the patients ^3^. Despite its effectiveness, there are patients intrinsically resistant to cisplatin-based therapies and an important fraction of tumors eventually develop chemoresistance ^4^

Cisplatin is composed of a double charged platinum ion surrounded by four ligands, two amines and two chlorides. Inside cells, the low chloride concentration facilitates cisplatin aquation replacing chloride groups by water molecules. This process produces a hydrolyzed (or aquated) form of cisplatin that is a potent electrophile (attracted to electrons) that can react with any nucleophile, including nucleic acids and the sulfhydryl groups of proteins ^5^.

Cellular cisplatin activity is a double-edged sword. In the nuclei, cisplatin produces DNA intra-and interstrand crosslinks that lead to apoptosis^5^. In the cytoplasm, due to its electrophilic activity, cisplatin behaves as an oxidant (loss of electrons results in oxidation) binding to proteins, including mitochondrial proteins, and especially to thiol-groups (-SH). Thus, cisplatin produces a ROS-homeostasis imbalance leading to more oxidizing conditions, which violates normal cellular function and ultimately can also promote apoptosis ^6^. Due to its broad and unspecific mode of action, cisplatin also affects normal cells. Nephrotoxicity, neurotoxicity and ototoxicity are some of the dose-limiting side effects reported upon cisplatin therapy ^5^. Nevertheless, the major obstacle for the clinical efficacy of cisplatin as an anticancer drug is the chemoresistance developed by tumors rather than its toxicity in normal cells.

Cisplatin resistance acquisition is multifactorial. The mechanisms by which tumor cells become resistant to the action of cisplatin have been classified in three types ^7^: (i) Pre-target mechanisms; reducing intracellular accumulation of cisplatin or increasing sequestration of cisplatin by nucleophilic scavengers as glutathione (GSH), metallothioneins, and other cysteine rich proteins, (ii) On-target mechanisms; acquiring the ability to repair adducts or becoming tolerant to unrepaired DNA lesions; and (iii) Post-target mechanisms; hampering the execution of apoptosis in response to DNA damage ^8^.

*C. elegans* is a well-established organism to study signaling pathways in response to drug exposure ^9^. Previous studies, performed in distinct biological contexts ^10, 11^ have revealed the value of *C. elegans* to identify genes related to the cisplatin response (**Supplementary Table 1**). Here we show for the first time that cisplatin in *C. elegans* produces DNA adducts and a systemic response driven by two conserved transcription factors. We have uncovered a sperm-specific sensitivity to cisplatin and have demonstrated that resistance of the nematode to cisplatin relies on the presence of a single evolutionary conserved aminoacid. Thus, this report is a more comprehensive study of the global response of *C. elegans* to cisplatin that also establish a reliable methodology to future studies on mechanisms of resistance to cisplatin that would continue contributing to the search of novels targets and makers for the benefit of cisplatin-based therapies.

## RESULTS

### A reliable assay to study the effect of cisplatin in *C. elegans*

In *Caenorhabditis elegans* nematodes cisplatin produces a wide variety of phenotypes depending on the concentration, length of treatment or developmental stage of the treated animals (**Supplementary Tables 1 and 2**). To implement a reliable methodology and systematically investigate the response of *C. elegans* to cisplatin, we established dose-response patterns during larval development, where cell divisions occur in somatic and germ cells. We exposed a synchronized population of L1 larvae on NGM plates to different cisplatin concentrations for 96 hours. We observed effects ranging from a developmental delay at 50μg/ml to a larval arrest at 200μg/ml (**Fig. 1a**). Based on this assay we concluded that body length at 48 post-L1, upon cisplatin exposure from 50 to 75μg/ml, is a reliable indicator of the cisplatin effect during *C. elegans* development **(Fig. 1b)**. Thus, we established a methodology to investigate how distinct treatments or gene activities can influence the response of *C. elegans* to cisplatin.

**Figure 1.**
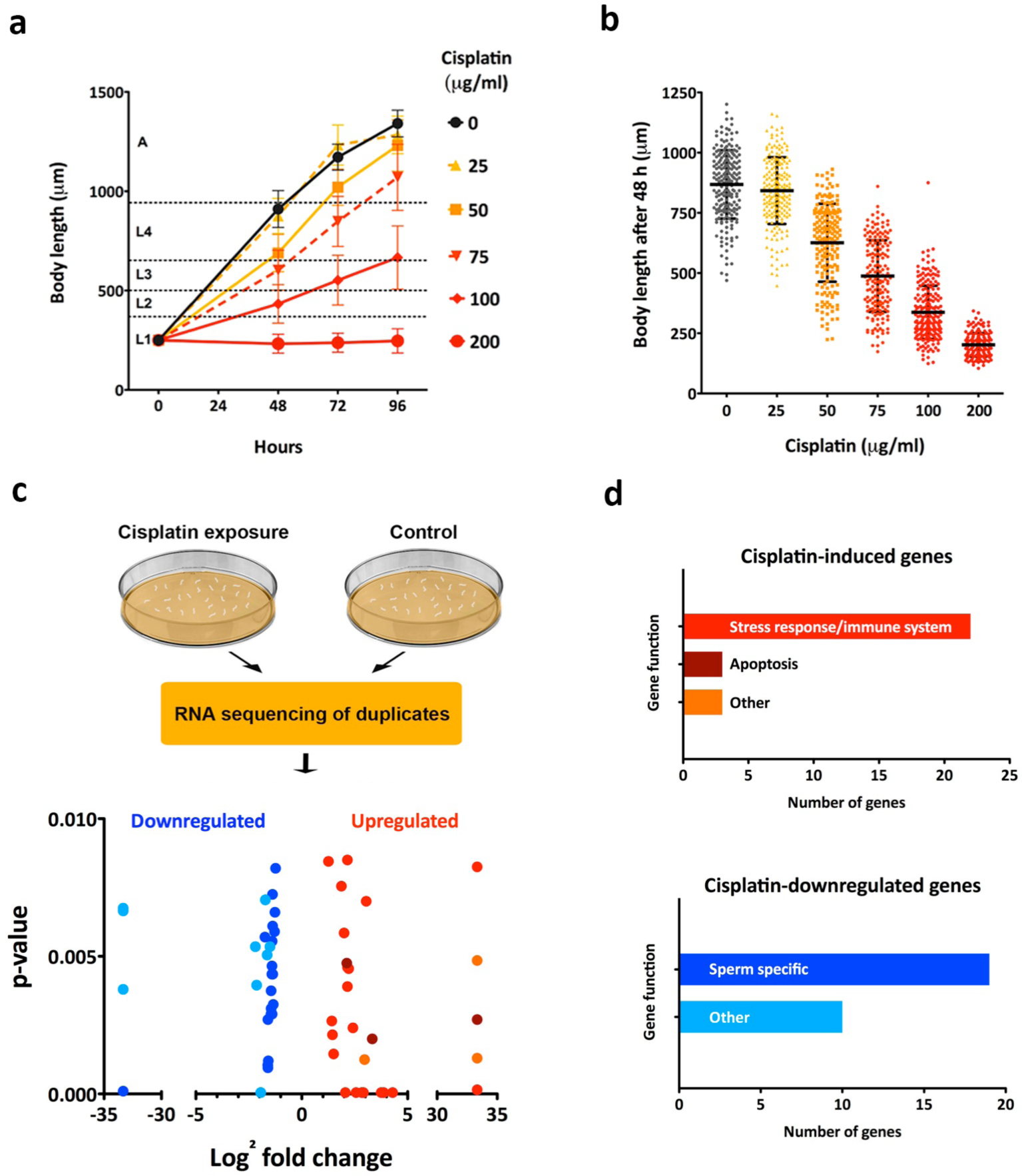
Dose-response effect of cisplatin in *C. elegans* development and RNA-seq of animals exposed to cisplatin. (A) Effect of distinct concentrations of cisplatin on larval development. Synchronized L1 larvae were grown on agar plates and exposed to different doses of cisplatin for 96 hours at 20°C. Body length values represent mean and SD (n=50) of two different experiments. (B) Body length of nematodes that were grown from L1 stage exposed to different doses of cisplatin for 48 hours at 20°C. Bars represent mean and SD of three independent replicates (n=50). (C) Schematic representation of transcriptomic analyses. Wild-type *C. elegans* from mixed stages were cultured with and without cisplatin (60μg/ml) for 24 hours at 20°C. Then, total mRNA of two biological replicates were purified and sequenced. Volcano plot represents genes significantly up-and down-regulated (*p*<0,01) in cisplatin-treated versus untreated animals. (D) Color code represents the functional categories of up-or down-regulated genes. Genes were clustered according to the gene functional classification tool DAVID and GO terms retrieved from Wormbase.

### Transcriptional response of *C. elegans* to cisplatin

To explore the global response of nematodes to cisplatin we studied transcriptional signatures in RNA-seq experiments. We performed a transcriptome analysis of a mixed-stage nematode population that was treated for 24 hours with 60 μg/ml of cisplatin since this dosage of cisplatin produces some phenotypes, but does not compromise the viability of the animals. To reduce the number of false positives, we used two biological replicates to compare the transcriptomes of treated and non-treated populations (**Fig. 1c**). We used the Cufflinks algorithm ^12^ to process the RNA sequencing data and study differential expression of genes. We identified a set of 83 genes up-regulated and 78 genes down-regulated by cisplatin in both experiments (*p≤0,05*) (**Supplementary Table 3**). A more restricted list of candidates, using a cut-off p-value of ≤0,01, includes 28 genes up-regulated and 29 down-regulated upon cisplatin exposure.

To further explore the functions of these 57 genes we performed a gene ontology (GO) analysis and identified predicted protein domains (COG) ^17^ (**Fig. 1d**). Genes encoding CUB-like domains, C-type lectins and Glutathione-S-transferases were found among the genes up-regulated upon cisplatin exposure. In *C. elegans,* the expression of such genes is associated with detoxification, redox balance, stress response and innate immune system and it is often regulated by the transcription factors DAF-16 and SKN-1 ^15–18^(**Supplementary Table 4**). Moreover, as expected from the cisplatin capacity to induce apoptosis, genes involved in the apoptotic signaling cascade, including *egl-1* and *ced-13,* were found among the cisplatin-induced genes^19^. However, as described below, these two genes functions in a distinct manner upon cisplatin exposure.

Of the 29 genes down-regulated by cisplatin, we found that most of those were sperm specific genes ^20^, with the Major Sperm Protein (MSP) domain as the most frequent GO annotation (**Supplementary Table 5**). Therefore, the reduced brood size induced by cisplatin may be provoked, partly at least, by a specific effect of this agent in spermatogenesis and/or in sperm activity.

### Cisplatin reduces the germ cell pool and affect sperm functionality

Our transcriptome analyses indicated that male germline genes are particularly downregulated in the presence of cisplatin. To investigate the effect of cisplatin in the germline, we exposed L4 animals to cisplatin for 24 hours, because the switch from spermatogenesis to oogenesis occurs at this stage and therefore both processes can be affected ^21^. We observed a dose-dependent reduction of the brood size upon cisplatin exposure (**Fig. 2a**). Such reduction correlates with an increase in the number of unfertilized eggs laid (**Fig. 2b**) and decrease in the mitotic pool of the germline (**Fig. 2c**). Moreover, we observed that cisplatin-treated germlines displayed fewer mitotic nuclei that were increased in size (**Fig. 2d**), which is an effect described in germlines exposed to DNA damaging agents due to cell cycle arrest upon activation of the S-phase checkpoint ^22^.

**Figure 2.**
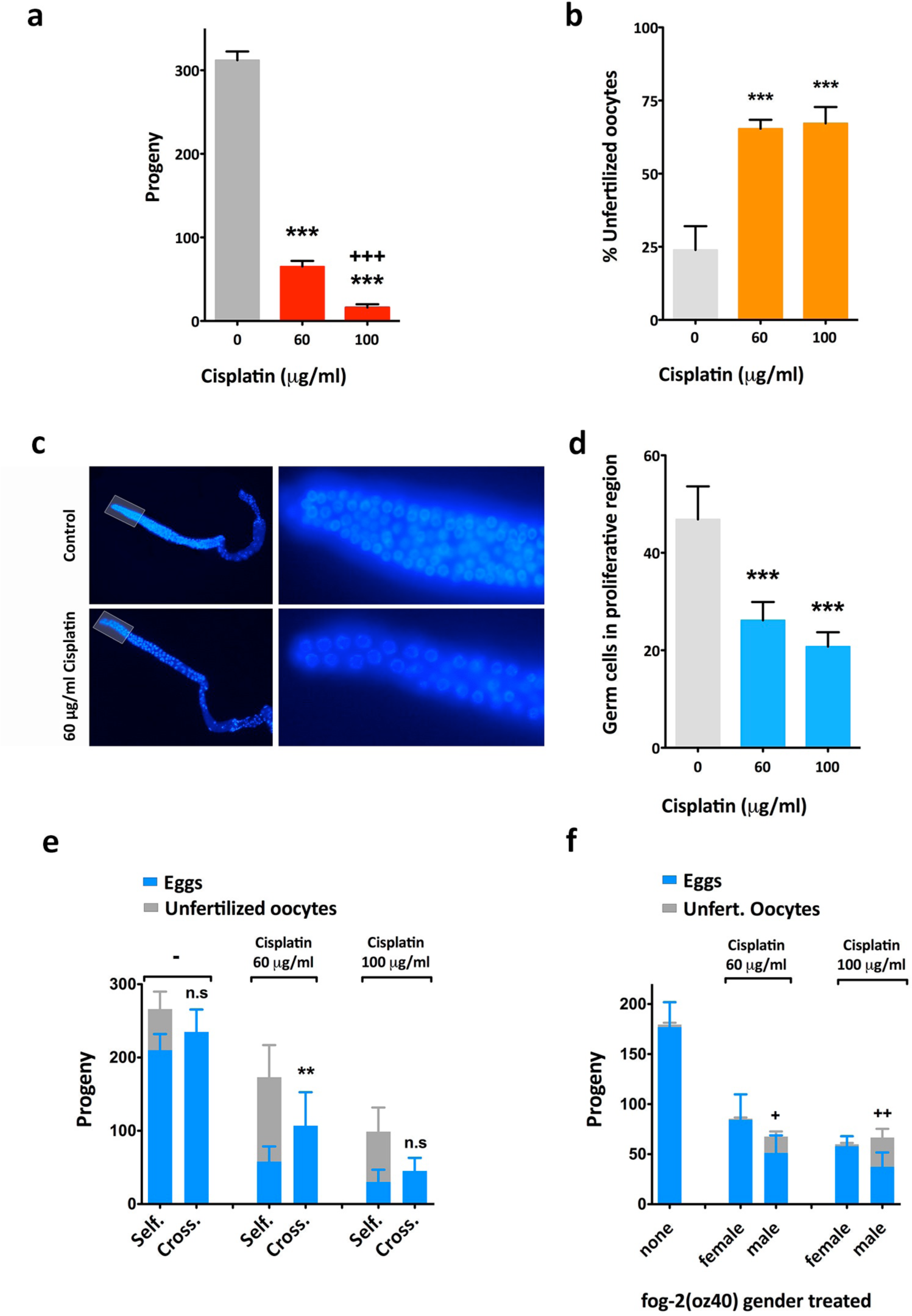
Effect of cisplatin in the germline. (A) Cisplatin reduces brood size. Progeny of *C. elegans* exposed to cisplatin at L4 stage for 24 hours at the indicated concentrations. (n=12, N=2). (B) Cisplatin increases the percentage of unfertilized oocytes laid. (n=12, N=2). (C) Representative DAPI staining of young adult germline exposed to cisplatin for 24h. Right panels show magnification of the germline proliferative region (highlighted area). (D) Cisplatin reduces the number of germ cells in the proliferative region of *C. elegans* germline. Number of cells corresponds to nuclei observed, in a single Z stack, at the proliferative region (50μm away from the distal end of the gonad). (n=15, N=2) (E) Cisplatin impact on sperm functionality. Cisplatin produces unfertilized oocytes and causes reduced brood size in self-fertilized/hermaphrodites worms “Self”, but this effect is partially rescued by crossing cisplatin-treated hermaphrodites with untreated males “Cross.”. (n=12, N=2). (F) Cisplatin-induced unfertilized oocytes are caused by defective sperm. *fog-2(oz40)* mutants do not produce sperm and mating is necessary for fertilization: “none” indicates that none of the sexes was treated; “female” indicates cisplatin-treated females fertilized by untreated males; and “male” indicates untreated females fertilized by cisplatin-treated males. Cisplatin produces unfertilized oocytes only when males are treated. (n=12, N=2). *++p>0,01* compared to female-treatment under same conditions. In all the graphs of this figure, bars show mean and SEM and Student’s t-test was applied. *\*\*\*p>0,001, **p>0,01, *p>0,1*

The increased number of unfertilized oocytes is a phenotypic hallmark of sperm failure in *C. elegans.* To investigate if the unfertilized oocytes observed upon cisplatin treatment (**Fig. 2e**) were the result of defective sperm, we crossed cisplatin-treated hermaphrodites with untreated males. We observed that sperm from untreated males rescued, partly at least, the brood size and abrogated the presence of unfertilized oocytes (**Fig. 2e**). These results suggest that the excess of unfertilized oocytes is due to the effect of cisplatin in spermatogenesis rather than in oogenesis. In other words, oocytes of cisplatin-treated animals are functional whereas sperm is not.

To further investigate the particular effect of cisplatin on sperm functionality we used *fog-2(oz40)*, which is a female/male strain instead of hermaphrodite. *fog-2(oz40)* females exposed to cisplatin and crossed with *fog-2(oz40)* untreated males showed a significant reduced brood size, as expected from the effect of cisplatin on mitotic cells, but we did not detect an increased production of unfertilized oocytes (**Fig. 2f**). On the other hand, by crossing *fog-2(oz40)* males exposed to cisplatin with untreated *fog-2(oz40)* females we observed a stronger reduction of the progeny, but also a higher number of unfertilized oocytes. These results confirm that, at similar dose, cisplatin does not hamper the capacity of oocytes to be fertilized, but affects the capability of sperm to fertilize.

### DAF-16 and SKN-1 are key players in the response to cisplatin

The abundance of dod (<u>d</u>ownstream-<u>o</u>f-<u>d</u>af-16) genes and other stress response genes in the list of genes upregulated upon cisplatin exposure led us to study the role of DAF-16 and SKN-1. These are the orthologs of human FOXO3a and Nrf2 respectively, and are required for stress resistance in *C. elegans* ^23^. Strikingly, we found that half of the up-regulated genes were previously reported as DAF-16 and/or SKN-1 targets (Fig. 3a).

**Figure 3.**
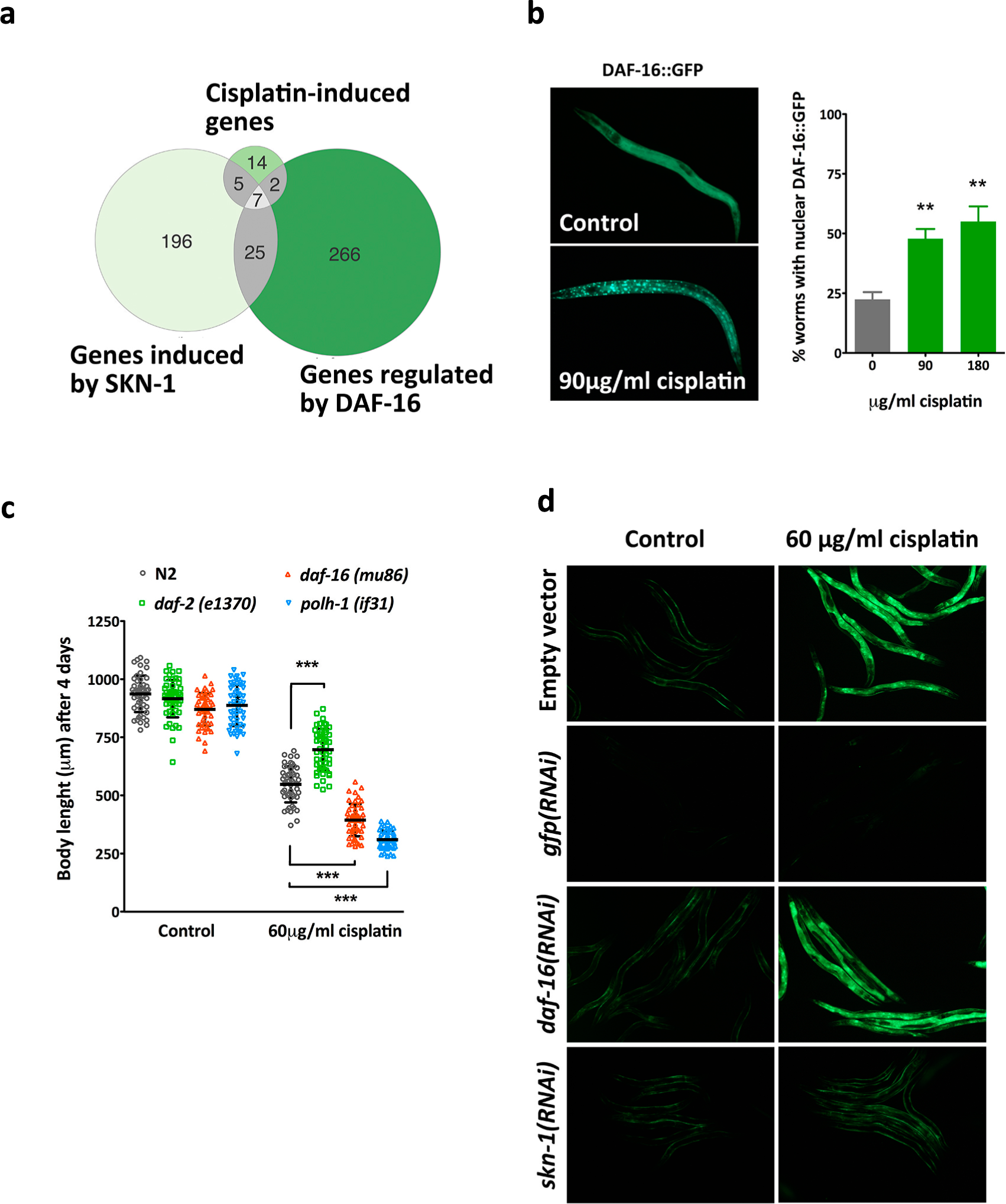
Role DAF-16/FOXO and SKN-1/Nrf2 transcription factors in the response to cisplatin. (A) Venn diagrams showing the significant overlap between the genes induced by cisplatin found in our transcriptomic analyses and genes regulated by DAF-16 ^18^ and SKN-1 ^65^. p<0,001 by Fisher’s test. (B) Cisplatin induces DAF-16 nuclear translocation. Representative image of L2 larvae carrying a translational GFP reporter of DAF-16 with and without cisplatin (90μg/ml for five hours). The graph represents the percentage of synchronized L2 larvae with DAF-16∷GFP predominantly nuclear after five hours of exposure to cisplatin at the indicated concentrations. Bars represent mean and SEM (n=50, N=3). ** p<0,01 relative to untreated worms by Student’s t-test. (C) IIS pathway influences the response to cisplatin. Body length of synchronized L1 larvae grown four days in the absence or presence of cisplatin. Grown at 15°C to avoid *daf-2(e1370)* dauer phenotypes. The inactivation of IIS pathway in *daf-2(e1370)* mutants, which keeps DAF-16 constitutively in the nucleus, causes resistance to cisplatin while *daf-16(mu86)* null mutant increases cisplatin sensitivity. Translesion synthesis polymerase 1 mutant allele *polh-1(if31)* was used as positive control ^66^. Bars represent mean and SD (n=50). *** *p<0,001* relative to untreated worms by Student’s t-test. (D) Cisplatin-induced activation of *gst-4* is regulated by SKN-1. Representative images of a synchronized *Pgst-4∷gfp* L4/young adults animals grown at 20°C on *daf-16(RNAi)* or *skn-1(RNAi)* bacteria from L1 stage in the presence or absence of cisplatin, *skn-1(RNAi)* inhibits the cisplatin-induced *gst-4* expression. Worms fed with *gfp* RNAi were used as positive control.

The evolutionary conserved insulin/insulin-like growth factor signaling (IIS) pathway, through its main transcription factor DAF-16/FOXO, controls many different biological processes and regulates a wide variety of stresses including starvation, oxidative stress ^24^, heavy metal toxicity ^25^ or ultraviolet radiation ^26^. DAF-16 is constitutively expressed and sequestered in the cytoplasm in its phosphorylated form. If the inhibitory phosphorylation is compromised, by e.g. reduced signaling of the DAF-2 Receptor-Tyrosin Kinase or stress conditions, DAF-16 will translocate to the nucleus. Using a DAF-16∷GFP-reporter strain we observed that DAF-16 nuclear location increased upon cisplatin exposure in a dose-dependent manner (**Fig. 3b**), confirming the implication of the IIS pathway in the response to cisplatin.

Next we investigated to what extent the manipulation of the IIS activity could modify the response of the organism to cisplatin. We observed that *daf-2(e1370)* mutants, which are animals lacking a functional DAF-2/IGF-1-like receptor that constitutively induce DAF-16 nuclear localization ^17^, display resistance to cisplatin during larval development (**Fig. 3c**). Consistently, *daf-16(mu86)* mutants were hypersensitive to cisplatin (**Fig. 3c**), highlighting the relevance of DAF-16 nuclear activity in the cellular response to cisplatin.

Differently from *daf-2* and *daf-16, skn-1* is an essential gene hampering the use of loss-of-function mutants. To confirm the implication of SKN-1 in the response to cisplatin we used a GFP reporter for one of its canonical targets, *gst-4.* SKN-1 regulates the stress-induced *gst-4* transcription in the presence of redox active compounds like paraquat or heavy metals ^27, 28^. In our RNA-seq data, *gst-4* is one of the SKN-1 regulated genes strongly induced by cisplatin and we validated this result using a *gst-4* transcriptional GFP reporter (**Fig. 3d**). Moreover, using *skn-1* RNAi we confirmed that the cisplatin induction of *gst-4* expression was *skn-1* dependent and *daf-16* independent (**Fig. 3d**). Thus, by studying the impact of cisplatin in the global gene expression we uncovered a systemic response of the organism that is driven by two conserved transcription factors. The fact that these transcription factors are effectors of metabolic and environmental signals opens new avenues to regulate the response to cisplatin in multicellular organisms.

### The BH3-only protein CED-13 protects against Cisplatin

We demonstrated that cisplatin leads to the formation of DNA adducts in *C. elegans* (**Fig. 4a**) and induces the expression of two-apoptosis related genes, *egl-1* and *ced-13,* which encode BH3-only domain proteins

**Figure 4.**
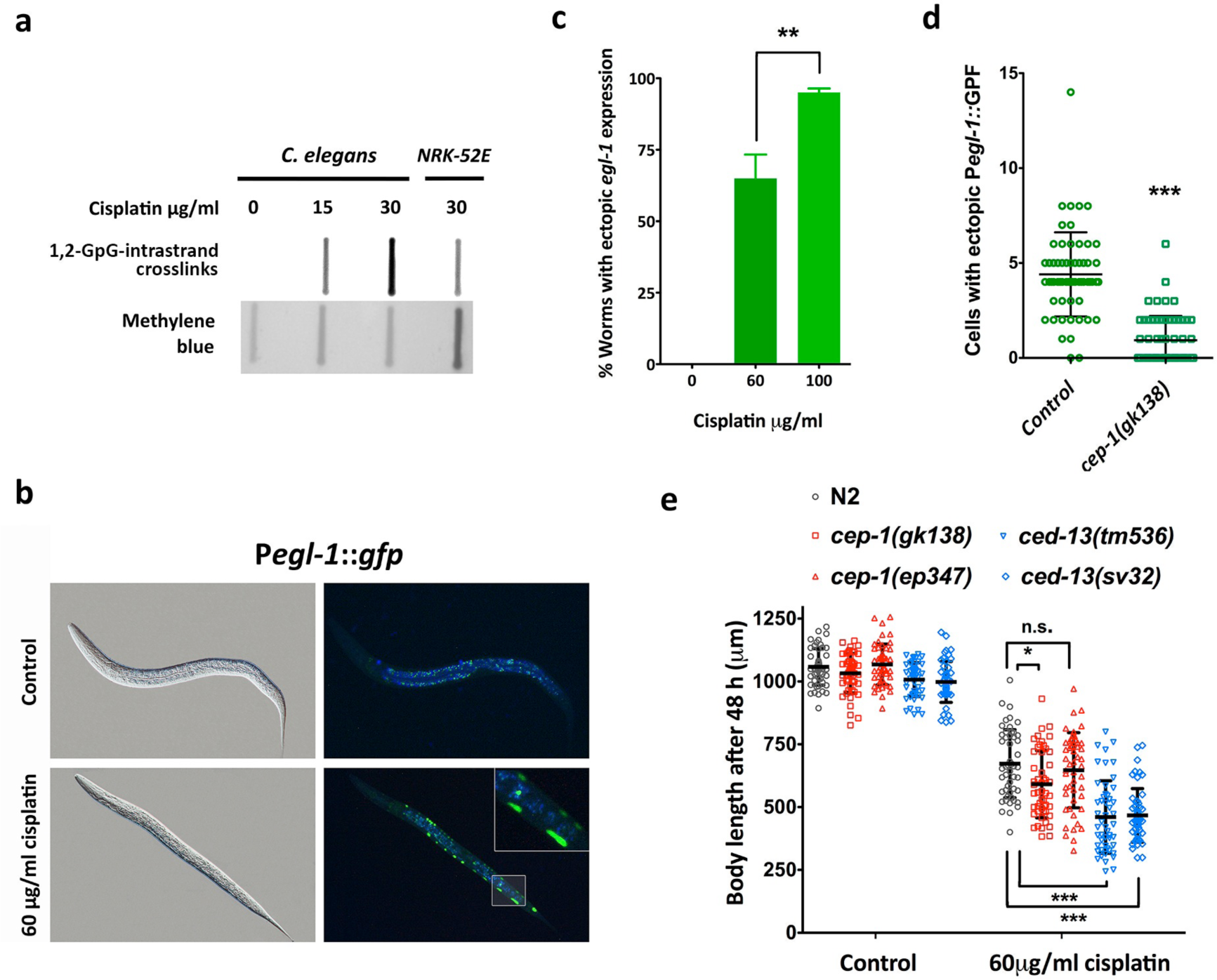
Role of apoptotic pathways in the *C. elegans* response to cisplatin. (A) Southern blot showing that 1,2-GpG-intrastrand crosslinks are raised upon cisplatin exposure of a *C. elegans* population. There is a dose-dependent accumulation of DNA adducts in presence of cisplatin. NRK-52E rat cells incubated with cisplatin were used as a positive control. (B) Cisplatin induces *egl-1* ectopic expression in somatic tissues during larval development. Representative images of L2 worms carrying a *Pegl-1∷gfp* reporter untreated and treated with cisplatin (60μg/ml for 24h at 20°C). (C) Quantification of worms with ectopic *Pegl-1*∷GFP fluorescence in somatic cells of synchronized L1 larvae grown at the indicated concentrations of cisplatin for 24h at 20°C. Bars represent mean and SEM of three different experiments (n=50). ** *p<0,01* by Student’s t test. (D) *egl-1* ectopic induction in somatic cells requires cep-1/p53. Quantification of cells with ectopic *Pegl-1*∷GFP fluorescence of L1 larvae exposed to 60 μg/ml of cisplatin for 24h at 20°C in wild type and *cep-1(gk138)* mutants. Bars represent mean and SEM of three different experiments (n=50) *** *p<0,01* by Student’s t-test. (E) Blockage of *cep-1/*p53 apoptotic pathway does not increase cisplatin resistance. Body length of L1 larvae of wild-type, and *cep-1* and *ced-13* null mutant alleles were grown at indicated concentration of cisplatin for 48 hours at 20°C. Bars represent mean and SD (n=50). This experiment was performed in triplicate with same results. *** *p<0,001* * *p<0,05* relative to wild type worms under same conditions by Student’s t-test.

*egl-1* and *ced-13* are transcriptionally induced upon DNA damage (also in case of UV or IR) and such induction is CEP-1/P53 dependent ^19^. *Y47G7B.2,* another cisplatin-induced-gene in our RNA-seq, encodes a nematode specific gene that is also upregulated upon DNA damage in a *cep-1* dependent manner ^19, 29^. Thus, the effects of cisplatin in upregulation of apoptotic genes could be due to *cep-1* dependent DNA damage.

*egl-1* is required for DNA-damage induced apoptosis in both somatic tissues and germline ^30^. We investigated *egl-1* induction in the soma using a GFP reporter strain (P*egl-1* ∷GFP). During larval development only 18 somatic cells of wild-type nematodes undergo apoptosis during early L2 ^31^. Strikingly, a 24 hour cisplatin-treatment in *C. elegans* (from L1 to L2) induces ectopic *Pegl-1* ∷GFP expression that was evident in somatic cells (**Fig. 4b, c**). Moreover, this cisplatin-induced *egl-1* ectopic expression was, at least partially, *cep-1* dependent (**Fig. 4d**).

In the canonical DNA-damage induced apoptotic pathway, *cep-1* is upstream of the BH3-only proteins EGL-1 and CED-13. However, *ced-13* has an additional role in the Intrinsic Apoptotic pathway that promotes protective changes rather than killing damaged irreparable cells ^32^. Accordingly, we found that *ced-13* mutants were sensitive to cisplatin (**Fig. 4e**) whereas two different *cep-1* mutants were not resistant to cisplatin (**Fig. 4e**).

In summary, during *C. elegans* larval development, cisplatin induces the expression of *egl-1* and *ced-13* but their influence is distinct. Inhibition of the DNA-damage induced apoptotic pathway by using *cep-1* and *egl-1* mutant does not produce animals resistant to cisplatin. On the contrary, *ced-13* mutants are sensitive to cisplatin underscoring a protective role of *ced-13* upon cisplatin exposure.

### A single selenocysteine confers systemic sensitivity to cisplatin

The majority of *C. elegans* genes related to cisplatin response published to date present cisplatin sensitivity if inactivated by RNAi or mutation (**Suplementary Table 1**). The human orthologs of these genes are potential targets of therapies to re-sensitize cisplatin resistant tumors. Given the clinical importance of identifying refractory patients, we wondered if our system was also valuable to study genes whose inactivation confers resistance to cisplatin highly relevant in the case to identify refractory patients. In the clinic, mutations or SNPs in these genes could function as predictive markers of cisplatin response. In human cell lines, thioredoxin reductase 1 (TRXR1) selenocysteine amino acid is a direct target of cisplatin producing cytotoxic and highly pro-oxidant TRXR1 forms called SecTRAPs, that lead to high redox stress resulting in cell death ^33^. TRXR1 is one of the few selenoproteins in mammals but TRXR-1 is the only selenoprotein (protein that includes a selenocysteine (SeCys) amino acid) in *C. elegans* ^34^ and, in contrast to mammals, this protein is not essential for viability ^35^. Inhibition of *C. elegans trxr-1* does not produce phenotypes in terms of morphology, growth, lifespan, brood size or response to oxidative stress. It is only essential for larval molting when *gsr-1* is inactivated in parallel ^35^. All these features make *C. elegans* a great multicellular system to specifically study the role of the SeCys residue in the response to cisplatin.

We tested the robustness of our model by studying the role of thioredoxin reductase 1 (*trxr-1*) in the *C. elegans* response to cisplatin. We used the CRISPR/Cas9 system to produce an endogenous TRXR-1 variant *trxr-1(cer5[U666STOP])* that changes the selenocysteine for a STOP codon (**Fig. 5a**). Similarly to the null mutant *trxr-1(sv47)*, this nonsense mutation showed larval arrest, due to molting defects, in combination with *gsr-1(RNAi)* (**Fig. 5b**). Strikingly, both the null mutant and the nonsense mutation removing the selenocysteine produce resistance to cisplatin (**Fig. 5c**). Thus, both the developmental function and the capacity to promote cytotoxicity in presence of cisplatin rely on the selenocysteine residue of TRXR-1.

**Figure 5.**
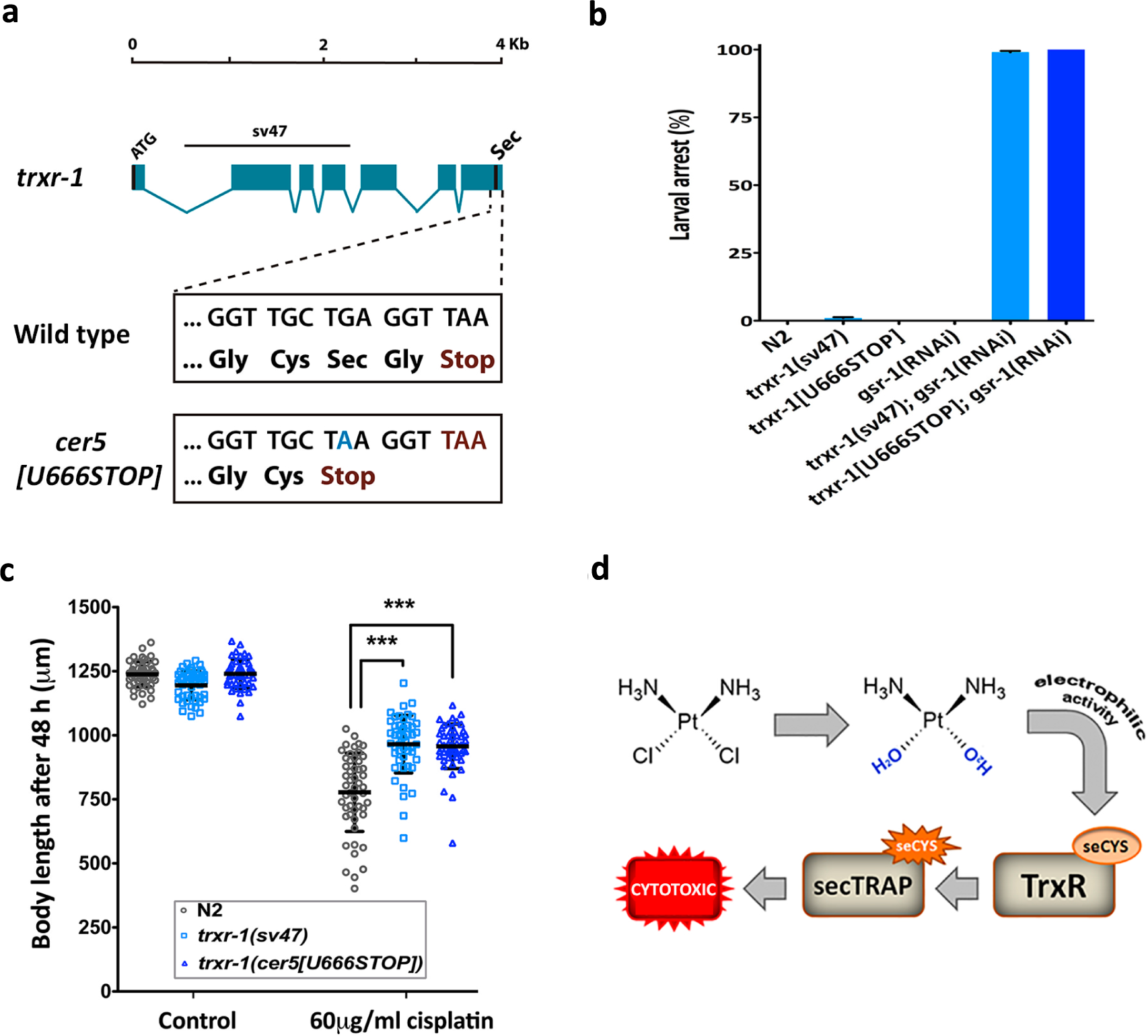
*trxr-1* null and nonsense mutants are resistant to cisplatin. (A) Representation of the *trxr-1* gene. Blue boxes represent exons, lines represent introns. The upper line denotes the 1663 bp region deleted in *sv47* allele. Selenocysteine codon ATG is the third last residue and is represented as Sec. The wild-type sequence of the last four codons and their corresponding aminoacids are boxed below. *cer5[U666STOP]* mutant allele edited by CRISPR/Cas9 carries a single substitution in Sec codon (A instead of G) generating a premature STOP codon that produces a Sec deficient TRXR-1 lacking the last two amino acids. (B) Sec deficient *cer5[U666STOP]* allele, similarly to *trxr-1* null allele, cause a molting-associated growth arrest when fed with *gsr-1(RNAi).* Graph shows the quantification of larval arrest of L1 larvae fed with *gsr-1(RNAi)* or control bacteria (empty vector). Bars represent mean and SD. (n=50, N=2). (C) *trxr-1(sv47)* null mutant and nonsense *cer5[U666STOP]* mutant strains show increased resistance to cisplatin. Body length quantification of synchronized L1 larvae grown on agar plates containing 0 and 60 μg/ml of cisplatin for 48 hours at 20°C. Bars represent mean and SD (n=50). This experiment was performed in triplicate with similar results. *** *p<0,001* relative to wild type worms under same conditions by Student’s t-test. (D) Squematic of the cytotoxic effect produced by the electrophilic activity of cisplatin when react with a selonocystein present in thioredoxin reductase 1.

Therefore, the electrophilic activity of hydrolyzed cisplatin in the presence of selenocysteine produce cytotoxic SecTRAPs forms that are major drivers of the harmful effect of cisplatin on multicellular organism (**Fig. 5d**).

## DISCUSSION

Despite three decades of clinical use and intense research, greater knowledge is necessary to understand, and ultimately control, the molecular events driving endogenous and acquired resistance to platinum-based chemotherapy ^36^. In this context, model organisms are platforms with growing interest in the study of the molecular, cellular and systemic responses to chemotherapeutic agents, but also to screen for new treatments ^37^. Here, we reinforce the use of *C. elegans* to study the complex response of multicellular organisms to cisplatin. The function of *C. elegans* genes in the response to cisplatin has been shown in different biological contexts as adult survival or germline apoptosis (Supplementary Table 1). After testing some of these contexts, we chose the body length of developmental larvae as a reliable and scalable system to evaluate the animal response to cisplatin. Nematodes are easy to synchronize at L1 stage and therefore we can study homogeneous populations during larval development where somatic and germ cells are in active proliferation and apoptotic pathways are functional.

The *C. elegans* germline is diverse in cell types since it contains mitotic cells, meiotic cells, mature oocytes and sperm. We observed that the effect of cisplatin in fertility and cell cycle arrest in the mitotic germ line is similar to that produced by other DNA-damaging insults as ionizing or ultraviolet C radiations ^22^, suggesting a direct action of cisplatin on DNA that is supported by our South-Western Blots for DNA adducts (1,2-GpG-intrastrand crosslinks). This cisplatin-induced DNA damage response (DDR) in the *C. elegans* germ line has already been documented ^38–40^. In addition, however, we found that defective sperm is a major cause of the reduced *C. elegans* fertility observed upon cisplatin exposure. Our results clearly show that sperm and oocytes are different in terms of resistance to cisplatin. Interestingly, cisplatin is particularly efficient for testicular tumors and cause a drastic effect on spermatogenesis and sperm in treated-patients, but such harmful effect seems to affect primarily the chromatin and is reversible ^41^. In females with ovarian cancer cisplatin also affect gametogenesis producing a reduction of the ovarian reserve ^42^. Detailed experiments need to be performed to better understand at which step the gametogenesis is hampered in the presence of cisplatin. In this sense, the hermaphrodite condition of *C. elegans* is a clear advantage to study the effect of any given cisplatin dose on oogenesis and spermatogenesis in the same organism.

Our RNA-seq analyses reveal a transcriptomic signature upon presence of cisplatin. The cisplatin-responsive genes identified could be involved in cell autonomous mechanisms (cells have an individual response to cisplatin uptake), or in non-cell autonomous mechanisms (the effect in certain cells influence distant cells). Since DNA is damaged by cisplatin, a cell-autonomous response occurs as checkpoint mechanism to favor DNA repair ^22^. On the other hand, the vast majority of genes induced by cisplatin in our experiments are related to general stress responses and the innate immune system, which suggests a coordinated systemic response. According to the genes deregulated, such response has DAF-16 and SKN-1 as major drivers and has similarities to that obtained from nematodes exposed to IR or UV-C ^29, 43^. DAF-16 activation positively regulates a wider spectrum of processes such as stress resistance, innate immunity or metabolic adaptation ^15, 44^ whereas the Nrf2 *C. elegans* ortholog SKN-1 is an important factor in detoxification and response to oxidative stress, and its downregulation has been associated to hypersensitivity to different stressor agents ^45^.

Interestingly, immune and stress reactions upon DNA damage are common in mammals ^46^, and similarly to what we observe in *C. elegans,* cisplatin induces the nuclear translocation of the DAF-16 homolog FOXO3A ^47^. Human FOXO proteins are crucial regulators of a multitude of cellular functions^48^ but our results encourage to further explore the manipulation of the IIS pathway to control the cellular response to cisplatin.

The SKN-1 human ortholog Nrf2 regulates the expression of stress responsive genes such as SOD, catalases or phase II detoxification enzymes like glutathione transferases ^49^. Nrf2 overexpression or hyperactivation provokes a direct effect in acquisition of resistance to a wide-spectrum of anticancer drugs in many cancer types ^49^–^50^. Accordingly, increased nuclear Nrf2 expression has been shown in cisplatin resistant bladder human cancer samples ^51^. On the contrary, inhibition of Nrf2 sensitizes cisplatin-resistant A549 cells ^52^. These observations are concordant to our findings in *C. elegans* suggesting that direct or indirect targeting of Nrf2 should be explored for cisplatin-combined therapeutic intervention.

Our transcriptomic analyses uncovered the upregulation of *egl-1* and *ced-13,* which are related to apoptosis. Using a *Pegl-1∷gfp* reporter we found ectopic expression in somatic cells. The apoptotic pathway in postembryonic somatic cells is not well described, but in the canonical view of the DNA-damage induced apoptotic pathway expression of the effector EGL-1 marks cells destined to die ^53^ and *cep-1/p53* is upstream ^54^. However, *Pegl-1∷gfp* is still overexpressed in *cep-1/p53* mutants exposed to cisplatin, indicating that inhibition of the DNA-damage induced apoptotic pathway do not completely protect the animal from cisplatin toxicity. Similarly, blockade of apoptosis in adult worms through *cep-1/*p53 inhibition does not have any effect on cisplatin sensitivity ^10^. Thus, inhibition of the DNA-damage apoptotic pathway is not an efficient strategy for animals to hamper the global effect of cisplatin. Still, expression of *cep-1/*p53 seems to be important for the full response to cisplatin, and p53 is required for the cytotoxic effect of cisplatin in human glioblastoma cells ^55^.

Surprisingly, we found that *ced-13* upregulation may have a protective role since *ced-13* mutants are sensitive to cisplatin. This result fits with the role of *ced-13* in the mitochondrial intrinsic apoptotic pathway that extends longevity ^56^. It is possible that post-mitotic cells, which are irreplaceable, activate *ced-13* activity to protect themselves from cisplatin. The *ced-13* role may be dependent on the insult levels or vary in distinct cell types since overexpression of *ced-13* in the soma induces cell death of somatic cells that normally survive ^29^.

Thioredoxin system proteins are key players in many important cellular processes, including the maintenance of the redox homeostasis ^57, 58^. Regarding the influence of thioredoxins in the cellular response to cisplatin, in mammalian cells it has been shown that a delicate balance exists between cytotoxic effect of the direct interaction with cisplatin, and the protective effect maintaining the redox equilibrium ^33, 59^. Studies in cell lines suggest that cisplatin cytotoxicity is promoted by a direct interaction between cisplatin and selenocysteine amino acids leading to the consequent formation of SecTRAPS ^33^. We confirmed the cytotoxicity of secTRAPS in *C. elegans* by making a *trxr-1* mutant that demonstrate that C-ter single selenocysteine (Sec) amino acid is required for the cytotoxic effect of cisplatin in the whole animal.

The multifactorial nature of cisplatin resistance hampers the finding of a unique solution to overcome the resistance of tumors to cisplatin. However, there are many parallelisms between the response of *C. elegans* and mammals to a cisplatin treatment that would facilitate the study in nematodes of particular genetic, metabolic and environmental factors influencing the resistance to cisplatin. This information would be of great help to identify predictive biomarkers and investigate new drugs to have more effective and personalized cisplatin-based therapies.

## MATERIAL AND METHODS

### Nematode strains and general methods

*Caenorhabditis elegans* strains were cultured and maintained using standard procedures ^60^, ^61^. The N2 (Bristol) strain was used as wild-type in all experiments and the following alleles and transgenic strains were used in this study: BS553: *fog-2(oz40)V*, XF132: *polh-1(if31)III*, CB1370: *daf-2(e1370)III*, TJ1; *cep-1(gk138)I*, CE1255: *cep-1(ep347)I*, FX536: *ced-13(tm536)X*, MD792: *ced-13(sv32)X*, VB1414: *trxr-1(sv47)IV*, CER151 *trxr-1(cer*5[U666Stop])IV, CF1038: *daf-16(mu86)I*, TJ356: *zIs356 (Pdaf-16∷daf-16∷gfp)*, WS1973: *opIs56(Pegl-1∷gfp)*, CER192: *cep-1(gk138)I*; *opIs56(Pegl-1∷gfp)*, CL2166 *dvIs19[pAF15(Pgst-4∷gfp∷*NLS)].

### CRISPR/Cas9

To generate the *trxr-1(cer5[U666Stop])* point mutation allele, we targeted a site near the selenocysteine codon with CRISPR/Cas9. To make the sgRNA expression plasmid we cloned the annealed oligonucleotides into BsaI-digested U6∷sgRNA PMB70 ^62^. Repair template containing desired modifications were designed by Gibson Assembly Cloning Kit (New England BioLabs) and cloned into the pBSK plasmid (Addgene). Oligonucleotide sequences used to generate the 5’ and 3’ 1.5kb overlapping fragments are available under request. Both sgRNA and repair template were verified by sequencing using T4 and M13 primers respectively. Young adult wild type animals were injected with 30ng/μl sgRNA, 100ng/μl repair template, 30ng/μl of *Peft-3∷Cas9* vector ^63^ and 2.5ng/μl of *Pmyo-2∷tdtomato.* Single fluorescent progeny were isolated and screened for the presence of the mutation by PCR. We finally established and validated homozygous mutant lines by PCR and sequencing.

### RNA-sequencing analyses

A mixed population of worms representing all stages and growing under control conditions was exposed to 60μg/ml of cisplatin. After 24 hours at 20°C, cisplatin-treated worms and untreated control animals were washed with M9 buffer to remove bacteria and total RNA was extracted using TRIzol method. Total mRNA was subsequently enriched using mirVana miRNA isolation kit (Ambion) followed by poly-A capture. Library construction and Illumina’s HiSeq technology sequencing was performed following manufacturer instructions. Reads (100bp length) were processed and aligned using TopHat software to the *C. elegans* reference genome, version WBcel235.74, to produce BAM files. These BAM files were analyzed with the SeqSolve NGS software (Integromics, S.L.) using a false discovery rate of 0,05, and filtering reads displaying multiple mapping sites. SeqSolve uses Cufflinks and Cuffdiff ^12^ programs to perform differential gene expression analyses between samples *(p<0,005).* Expression values were normalized in FPKM (fragments per kilobase of exon per million fragments mapped). Datasets of up-and down-regulated genes was generated comparing the differential gene expression analyses of two independent replicates *(p<0,01* and *p<0,05).*

### Cisplatin assay during larval development

A synchronized population of L1-arrested larvae was cultured on nematode growth medium (NGM) plates containing fresh OP50 and 0-200μg/ml of cisplatin (Sigma). Body length of at least 50 worms for each condition was measured at 48, 72 and 96 hours at 20°C at the stereomicroscope using NIS-Elements 3.2 imaging system. The subsequent analyses were performed at 60μg/ml of cisplatin under same conditions except assays containing *daf-2(e1370)* strain, that were performed at 15°C measuring the body length after four days of incubation to avoid temperature-related developmental delay. Each assay was done in duplicate and at least two biological replicates were performed. Non-parametric Student t-test (GraphPad Prism 5) was used to determine the significance of differences in the mean.

### Analysis of cisplatin effect in the germline

To analyze the effect of cisplatin in the germline, a synchronized population of L4 state animals grown under standard conditions (39 hours at 20°C) was transferred onto NGM plates containing 60μg/ml cisplatin. After 24h of incubation at 20°C worms were washed in PBS and germ lines were dissected anesthetizing worms with 3mM of levamisol in PBS, fixed with 4% paraformaldehyde and stained with DAPI (0,6μg/ml). These stained gonads were photographed using a Nikon ECLIPSE TI-s inverted microscope. For germ cell quantification in the proliferative region, the total number of cells in a single Z stack within 50 mm of the distal end of the gonad was counted. At least 15 germ lines were counted for each experiment.

### Brood size and unfertilized oocytes laid assays

To perform the self-fertilization assays, a synchronized population of L3 stage worms was exposed to NGM plates adding fresh OP50 and 0, 60 or 100μg/ml of cisplatin for other 24 hours. A total of 12 worms of each condition were singled out onto fresh OP50 plates, counting the progeny and the number of oocytes laid after three days. Similar protocol was performed for crossing assays where we used either a male-enriched population in case of N2, or the male-female population of the *fog-2(oz40)* allele. After 24 hours of exposure to cisplatin, a total of 12 genetic crosses where performed for each condition, placing one hermaphrodite, or female in case of*fog-2(oz40)*, and four males in fresh 0P50 plates, letting them mate and lay progeny for three days. This experiment was performed in duplicate. Non-parametric Student t-test was used to determine the significance of differences in the mean.

### *In vivo* intracellular localization of DAF-16

To determine DAF-16 subcellular localization, a synchronized L2 stage population of worms carrying DAF-16∷GFP transgene (TJ356) were transfer to M9 buffer containing 0, 90 or 180μg/ml of cisplatin for five hours at 20°C. Then, worms were washed in M9 and mounted on a microscope slide containing 2% agar pad using a drop of levamisole 3mM to anesthetize them. We quantified the DAF-16 subcellular localization by considering as “nuclear” worms showing a mainly nuclear GFP accumulation along the whole body, as shown in figure 3b. A total of 50 worms were observed in each one of three independent replicates.

### *In vivo egl-1* and expression assay

To determine ectopic *egl-1* expression, a synchronized L1 stage population of *Pegl-1∷gfp* (WS1973) transgenic nematodes were grown in NGM plates containing fresh OP50 and 60 μg/ml cisplatin plates for 24 at 20°C. For *in vivo* observation, worms were recovered with M9 buffer, washed and mounted on a microscope slide with a 2% agar pad using a drop of 3mM of levamisole. We used a Nikon ECLIPSE TI-s inverted microscope to quantify the number of cells showing ectopic GFP signal (N=50). These experiments were performed three independent times.

### RNAi mediated interference and *in vivo gst-4* expression assay

*daf-16 and skn-1* RNAi clones used in this study were obtained from the ORFeome libray ^64^. RNAi by feeding was performed following standard conditions; using NGM plates supplemented with 50 μg/μl ampicillin, 12.5μg/μl tetracycline and 3mM IPTG. To analyze *gst-4* induction, a synchronized L1 stage population of *Pgst-4∷gfp* reporter strain was grown on plates containing the corresponding RNAi clone for 24 hours at 20°C. Then, worms were transferred to new RNAi plates including 0 or 60μg/ml cisplatin for 24 hours. Then, worms were recovered with M9 buffer, washed and mounted on a microscope slide containing 2% agar pad using a drop of 3mM of levamisole. Nikon *ECLIPSE TI-s* inverted microscope was employed for *in vivo* observation.

### Southwestern blot analysis for Pt-DNA adducts

The Southwestern blot analysis was performed using DNA extracted from a synchronized population of *C. elegans* after 24 h incubation with or without cisplatin. Genomic DNA (0.5 μg each sample) from *C. elegans* and from NRK-52E rat cells (positive control) was isolated using the “DNeasy Blood and Tissue” kit (Qiagen (Hilden, Germany)), denatured by heating (10 min, 95°C) and cooled on ice. After adding 100μ! ice-cold ammonium acetate (2M) the DNA was transferred onto a nitrocellulose membrane, which was soaked in 1M ammonium acetate before. After washing (1M ammonium acetate and water) the membrane was baked for 2 hours at 80°C before it was blocked in 5% non-fat milk in TBS/0.1% Tween 20 overnight at 4°C. Incubation with the primary antibody directed against Pt-DNA adducts (1:2000; abcam ab103261) was conducted for 1 h at RT. Visualization of the antibody signal was done by chemiluminescence (Biorad ChemiDoc Touch Imaging System). Additionally, the membrane was stained with methylene blue (MP Biomedicals (Santa Ana, CA, USA)) to ensure equal DNA loading.

### Genetic interaction with *gsr-1* assay

This procedure was performed as previously described ^35^. Five L4 worms of the corresponding genotype were transferred to plates containing *gsr-1(RNAi)* bacteria for 24 hours at 20°C. Then worms were transferred to new fresh *gsr-1(RNAi)* plates, allowing lying eggs for 12 hours and then removed. After 3 days at 20°C larval arrest was analyzed only in day-2 plates.

## SUPPLEMENTARY TABLES

**Table S1**. Previous studies on cisplatin in *C. elegans*

**Table S2. Phenotypes observed in *C. elegans* upon different cisplatin treatments.** Phenotypes are abbreviated as follows: *Rbs* (Reduced brood size), *Emb* (Embryonic lethality), *Pvu* (Protruding vulva), *Ste* (Sterile), *Lav* (Locomotor activity variant), *Let* (lethal), *Gro* (Growth delay), *Uol* (Unfertilized oocytes laid).

**Table S3. Excel file containing the list of genes deregulated upon ciplatin exposure.** Sheet 1: genes deregulated with a *p*-value<*0,05.* Sheet 2: genes deregulated with a *p*-value<*0,01*

Table S4. Genes induced by cisplatin. Genes whose expression was significantly increased in two different experiments after a 60μg/ml cisplatin treatment. *p*-value<*0,01.* Columns on the rights indicate if these genes are regulated by DAF-16 or SKN-1 according to published literature.

Table S5. Genes downregulated by cisplatin. Genes whose expression was significantly decreased in two different experiments upon a 60μg/ml cisplatin treatment. *p*-value<*0,01*.

